# Endocytosis of very low-density lipoprotein particles: an unexpected mechanism for lipid acquisition by breast cancer cells

**DOI:** 10.1101/684274

**Authors:** Leslie E. Lupien, Katarzyna Bloch, Jonas Dehairs, William W. Feng, Wilson L. Davis, Thea Dennis, Johannes V. Swinnen, Wendy A. Wells, Nicole C. Smits, Nancy B. Kuemmerle, William B. Kinlaw

## Abstract

We previously described the expression of CD36 and lipoprotein lipase (LPL) by breast cancer (BC) cells and tissues, and the growth-promoting effect of very low-density lipoprotein (VLDL) supplementation observed in BC cell lines only in the presence of LPL. We now describe the deployment of LPL by BC cells. Our data support a model in which LPL is bound to a heparin-like heparan sulfate proteoglycan motif on the BC cell surface and acts in concert with the VLDL receptor to rapidly internalize intact lipoproteins via receptor-mediated endocytosis. We further observe substantial alterations in gene expression programs related to pathways for lipid acquisition (synthesis vs. uptake) in response to each the availability of exogenous triglyceride in tissue culture media and LPL expression status. Current literature emphasizes *de novo* fatty acid synthesis as the paramount mechanism for lipid acquisition by cancer cells. Our findings indicate that exogenous lipid uptake can serve as an important method of lipid acquisition for cancer cells, alongside *de novo* lipogenesis, and that the relative reliance on these two modes of lipid acquisition may vary among different BC cell lines and in response to nutrient availability. This concept has obvious implications for the development of therapies aimed at the lipid dependence of many different cancer types. Moreover, the mechanism that we have elucidated provides a direct connection between dietary fat and tumor biology.

---

The dependence of several tumor cell types, including breast cancer, on a supply of fatty acids to maintain proliferation is well established [1]. The literature emphasizes *de novo* lipid synthesis as the paramount mechanism used by tumor cells to satisfy this dependency. *De novo* lipogenesis is a cytosolic process that produces the saturated long-chain FA palmitate (C16:0), the bulk of which is used for plasma membrane phospholipid synthesis. This, plus the low rates of FA synthesis observed in most nonmalignant tissues, has positioned FA synthetic enzymes, especially fatty acid synthase (FASN), as therapeutic targets, and has prompted efforts to develop small molecule FASN inhibitors, reviewed in Kinlaw, Baures [2].

In this context, we and others have observed that provision of exogenous FA to cancer cells permits evasion of the cytotoxic effects of FASN inhibition, revealing the ability of cancer cells to take up, as well as synthesize, lipids [3, 4]. We further demonstrated that BC cells and clinical BC tissues robustly express LPL, the principal enzyme for the hydrolysis of triglyceride (TG) carried in lipoproteins, as well as the cell surface channel for FA uptake, CD36. Others have begun to highlight the presence of LPL in and on the surface of cancer cells, as either an expressed protein or one acquired from adjacent LPL-secreting cells, such as adipocytes and macrophages [5-11]. The most prominent example is in chronic lymphocytic leukemia (CLL) where a large body of evidence supports *LPL* as one of the strongest prognostic biomarkers in the disease [12-16]. Efforts have been made to understand the molecular mechanisms regulating LPL expression [17-20], and the functional role of LPL, particularly in poor-prognosis IgHV-unmutated CLL. Despite many recent advances [19-24], this continues to be an active area of research, and the role of LPL in the metabolism and pathophysiology of cancer in general, and CLL in particular, is still largely unresolved.

LPL is known most prominently as the enzyme responsible for the hydrolysis of TG in circulating lipoproteins. Here, LPL is produced by myocytes and adipocytes and secreted into the surrounding interstitial spaces, where it forms a dynamic reservoir through transient interactions with heparan sulfate proteoglycans (HSPGs). From this location, it is transported to its site of action in the capillary lumen [25]. For years, the dogma held that secreted LPL was tethered to capillary endothelial cells solely by electrostatic interactions between its positively charged heparin-binding domains and the negatively charged HSPGs on the endothelial cell surface [26-29]. This belief was supported by the fact that LPL can be readily released into the plasma by heparin [30], and by countless *in vitro* studies showing that LPL binds to HSPGs and that this interaction can be disrupted by reducing the sulfation of HSPGs or by digesting them with heparinase and/or heparitinase treatment [31-33].

An alternate model has come to light in recent years involving glycosylphosphatidylinositol-anchored high-density lipoprotein-binding protein 1 (GPIHBP1). In this model, LPL is expressed by parenchymal cells, secreted into the interstitial spaces, captured by GPIHBP1 on the antiluminal surface of local capillary endothelial cells, and then shuttled to the luminal surface of the capillary [25, 34, 35]. Here, GPIHBP1 facilitates LPL binding to the luminal surface (“margination”) and acts as an anchoring site for LPL on the capillary wall, creating a stable “platform for lipolysis.” Once lipoproteins are bound to this platform, LPL mediates TG hydrolysis, releasing glycerol and free fatty acids (FFAs) that can then be taken up through channels, such as CD36, on the surface of underlying adipocytes and myocytes. An extensive body of evidence highlights the importance of LPL and GPIHBP1 in the maintenance of plasma TG levels [25, 35-40]. There are, however, many contexts in which LPL functions without association to GPIHBP1, including its role as a non-catalytic mediator of lipid uptake.

Apart from its lipolytic activity in the capillary, LPL has been shown to act as a noncatalytic bridge, promoting the accumulation and active uptake of lipoproteins and lipoprotein remnants [41] via receptor-mediated endocytosis [42]. In this role, LPL interacts with lipoproteins and a variety of different cell surface proteins, including HSPGs and members of the LDL receptor family, including the VLDL receptor (VLDLR) [43]. The ability of LPL to serve as a molecular bridge has been supported by both *in vitro* and *in vivo* experiments, including the work of Merkel et al., which showed that catalytically inactive LPL expressed in muscle could still bind to HSPGs and induce the uptake of VLDL [41, 44].

Our previous work described the expression of CD36 and LPL by BC cells and tissues, and the growth-promoting effect of VLDL supplementation observed in BC cell lines only in the presence of LPL. We now describe the deployment of LPL by BC cells. Our data support a model in which LPL is bound to a heparin-like HSPG motif on the BC cell surface and acts in concert with the VLDLR to rapidly internalize intact lipoproteins via receptor-mediated endocytosis. We further observe substantial alterations in patterns of gene expression related to pathways for lipid acquisition (synthesis vs. uptake) in response to the availability of exogenous TG in tissue culture (TC) media and/or LPL expression status. These findings highlight the importance of exogenous lipid uptake as a method of lipid acquisition for cancer cells, alongside *de novo* lipogenesis, and provide evidence for BC cell metabolic plasticity in response to nutrient availability.

## RESULTS

### LPL is in and on the surface of cancer cells and clinical breast tumor tissue. BC cells that do not express LPL may acquire it from exogenous sources, such as the FBS component of tissue culture media

Breast tumor tissues stained with an antibody directed against LPL displayed both cytoplasmic and cell surface LPL staining (**Figure 1A**). Using qRT-PCR, we established that LPL is variably expressed across a panel of human cancer cell lines (Figure S1). We previously described LiSa-2 liposarcoma cells, HeLa cervical cancer cells, and DU4475 BC cells as high LPL-expressing and -secreting cell lines. Here, using flow cytometry with our AF-647-LPL antibody, we show that LPL is found to a variable degree on the surface of LPL-expressing cancer cells and at higher levels inside the cell, with the “total” LPL pool represented by the fixed/permeabilized (F/P) condition (**Figure 1B**). Cell surface-bound LPL was displaceable by heparin (500 µg/mL), as quantified by flow cytometry (**Figure 1C**) and immunocytochemistry using non-adherent DU4475 TNBC cells with a protocol designed to preserve cell surface HSPGs (**Figure 1D**).

**Figure 1.**
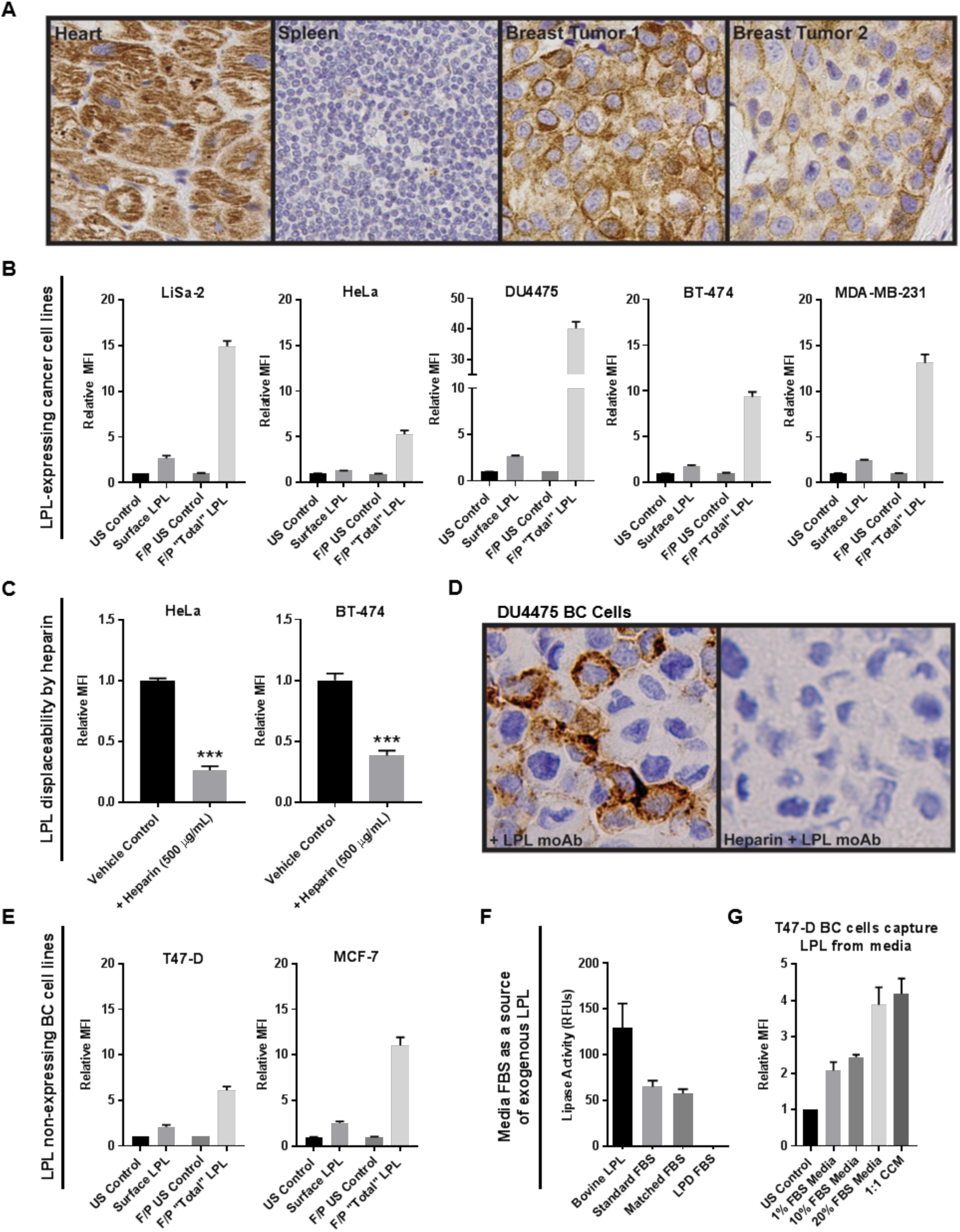
LPL is in- and on the surface of breast tumor cells and cancer cell lines and is displaceable by heparin. (**A**) Control (heart and spleen) and breast tumor slides were stained for LPL. Breast tumor cells display cell surface and cytoplasmic LPL staining. (**B**) The presence of LPL protein in- and on the surface of cancer cells was assessed via flow cytometry. For all cell lines, the “total” LPL present in fixed/permeabilized (F/P) cells exceeded that on the cell surface. Note different scales; error is SEM of > 3 experiments. (**C**) Representative data from heparin displacement studies using LPL-expressing HeLa cervical cancer and BT-474 BC cells. Heparin significantly displaces LPL from the cell surface, as detected by flow cytometry (***p < 0.001, two-tailed unpaired t-test with Welch’s correction). (**D**) Immunocytochemical analysis of LPL on the surface of DU4475 non-adherent TNBC cells. Cells were incubated with LPL antibody ± heparin (40 µg/mL), embedded in agarose, formalin fixed, embedded in paraffin, and sectioned. Cell surface LPL antibody was detected with peroxidase conjugated anti-IgG (brown pigment) with hematoxylin counterstain (blue). (**E**) T47-D and MCF-7 BC cells had no detectable LPL expression in our qRT-PCR studies. However, LPL was found on the cell surface and, to a greater extent, inside these cells by flow cytometry. (**F**) Standard, LPD, and matched control FBS were assessed for lipase activity. Activity was detected in bovine LPL and in standard and matched control FBS, but not LPD FBS. (**G**) T47-D BC cells were incubated in different media overnight to directly investigate whether BC cells can capture LPL from media. A FBS dose-dependent increase was observed in the presence of cell surface LPL. LPL was also detected in cells incubated with culture media conditioned by LPL-secreting LiSa-2 liposarcoma cells.

We predicted that LPL would be present in cells that express the gene but were surprised to find that LPL was detectable in and on the surface of T47-D and MCF-7 LPL low to non-expressing BC cell lines (**Figure 1E**). We hypothesized that without LPL mRNA expression, the protein must be acquired from an exogenous source, such as the FBS of the TC media. A lipase activity assay confirmed that detectable lipase activity was present in bovine LPL and in standard and matched control FBS, but not in lipoprotein-depleted FBS (**Figure 1F**). We assessed the ability of T47-D cells to capture LPL from the TC media by incubating cells in media harboring different amounts of FBS overnight (**Figure 1G**). A FBS concentration-dependent increase was observed in the LPL present on the surface of T47-D cells. LPL was also detected on cells incubated with media conditioned for three days by LPL-secreting LiSa-2 liposarcoma cells.

### The HSPG motif, HS^NS4F5^, is readily detectable on the surface of BC cells and serves as a binding site for LPL

LPL has been reported to bind to HSPGs, the (GlcNS6S-IdoA2S)_3_ or HS^NS4F5^ motif, in particular [33, 45]. Using qRT-PCR, we first determined that the cancer cell lines used in our studies did not express detectable levels of GPIHBP1, the one exception being DU4475 TNBC cells (not shown). Using flow cytometry and a Dylight 650-labeled antibody for HS^NS4F5^ (NS4F5^IgG^) we established that this binding site for LPL is present on the surface of cancer cells (Figure S2A). This was confirmed visually using fluorescence confocal microscopy, shown by a representative image of HS^NS4F5^ staining on the surface of MDA-MB-231 BC cells (Figure S2B). The abundance of HS^NS4F5^ on the cell surface was sensitive to several factors, including confluence and nutrient supply (not shown). Knockdown of heparan sulfate 6-O-sulfotransferase 1 (HS6ST1), an enzyme responsible for 6-O-sulfation of heparan sulfate, resulted in significant reduction of HS^NS4F5^ on the surface of MDA-MB-231 BC cells, supporting the specificity of the NS4F5^IgG^ (Figure S2C).

### BC cells take up intact VLDL particles in a temperature-, concentration- and time-dependent manner through receptor-mediated endocytosis

Our initial studies revealed that VLDL particles bind to- and are rapidly internalized by cancer cells. This raised the question of whether the observed uptake represented the internalization of intact lipoprotein particles or that of FFA released by LPL-mediated hydrolysis of TG. To investigate this, the lipid- and protein-components of VLDLs were labeled with DiI- and DyLight 650, respectively, and visualized using confocal microscopy (Figure S3). As shown in the merged channel, the fluorescently labeled lipid- and protein-components of internalized VLDL particles coincide, indicating that intact lipoproteins are internalized.

We characterized DiI-VLDL uptake using flow cytometry, confocal microscopy, and plate-based fluorescent assays. With MDA-MB-231 BC cells as our model, we show that DiI-VLDL uptake is time- and dose-dependent, and temperature-sensitive (**Figure 2**). MDA-MB-231 cells incubated at 4°C with DiI-labeled VLDL particles (5 µg/mL) bind VLDLs at the cell surface, while incubations at 37°C result in both binding and internalization of the lipoprotein particles (**Figure 2C**, **D**). Evidence that VLDL uptake involves the binding and rapid internalization of intact lipoprotein particles through temperature-dependent process led us to investigate whether VLDLs are taken up through receptor-mediated endocytosis. Treatment of MDA-MB-231 cells with dynasore, an inhibitor of GTPase-dependent clathrin-coated pit-mediated endocytosis, resulted in a concentration-dependent reduction in DiI-VLDL uptake, with concentrations widely used in the literature [46] reducing uptake to less than 1% of the control (**Figure 2E**).

**Figure 2.**
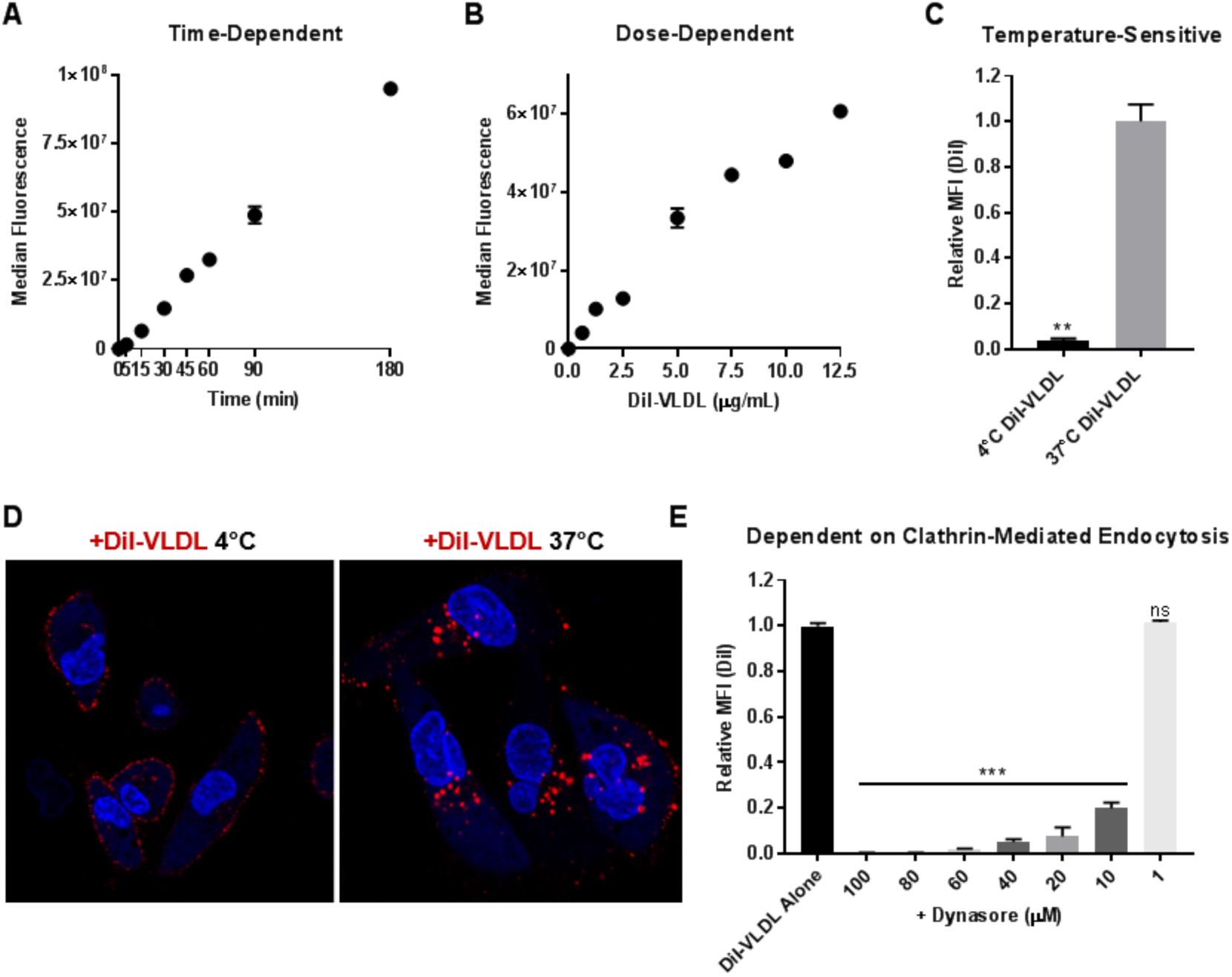
VLDL particles bind to- and are internalized by cancer cells. DiI-VLDL uptake is temperature-, dose-, and time-dependent, and is inhibited by treatment with the dynamin GTPase inhibitor, dynasore. Median fluorescence of MDA-MB-231 BC cells incubated with (**A**) DiI-VLDL (5 µg/mL) for durations of time ranging from 5 min to 3 h, and (**B**) DiI-VLDLs for 45 min at increasing dosage. (**C**-**D**) MDA-MB-231 BC cells were incubated with DiI-VLDL (5 µg/mL, 45 min) in 37°C or 4°C. (**C**) Cells at 37°C displayed significantly more uptake than those incubated at 4°C as quantified by flow cytometry (**p <0.01, two-tailed unpaired t-test with Welch’s correction). Duplicate or triplicate for all data points; error is SD. (**D**) Visualization via confocal microscopy shows that DiI-VLDLs remain bound at the cell surface at 4°C, whereas incubation at 37°C results in binding and internalization of VLDL particles. (**E**) Relative uptake of DiI-VLDL particles following treatment with dynasore (30 min at 37°C) measured by flow cytometry. Concentration-dependent reduction in DiI-VLDL uptake with dynasore treatment, ***p < 0.001, one-way ANOVA with correction for multiple comparisons.

### DiI-VLDL binding and uptake are abrogated by treatment with heparin, heparinase, or an antibody targeting the HS^NS4F5^ motif

We hypothesized that VLDL uptake was mediated by cell surface HSPG-bound LPL acting as a non-catalytic bridge to facilitate the uptake of intact lipoprotein particles through receptor-mediated endocytosis. We assessed the dependence on HSPGs (directly) and LPL (indirectly) by first testing the effect of heparin on DiI-VLDL uptake. Treatment with heparin resulted in a concentration-dependent decrease in the DiI-VLDL uptake by MDA-MB-231 cells (**Figure 3A**). Treatment with heparinase I and/or III enzymes (to digest cell surface HSPGs) likewise reduced DiI-VLDL uptake compared to the vehicle control (**Figure 3B**). Confocal microscopy using DiI-VLDLs and DyLight 650-NS4F5^IgG^ shows that both VLDLs and HS^NS4F5^ localize to the cell surface (in cells incubated at 4°C) (**Figure 3C**). Treating cells with unlabeled NS4F5 antibody reduced DiI-VLDL uptake, measured by flow cytometry (**Figure 3D**). Together, these data support a model in which HSPGs, particularly the HS^NS4F5^ motif, are integral for VLDL uptake by MDA-MB-231 BC cells.

**Figure 3.**
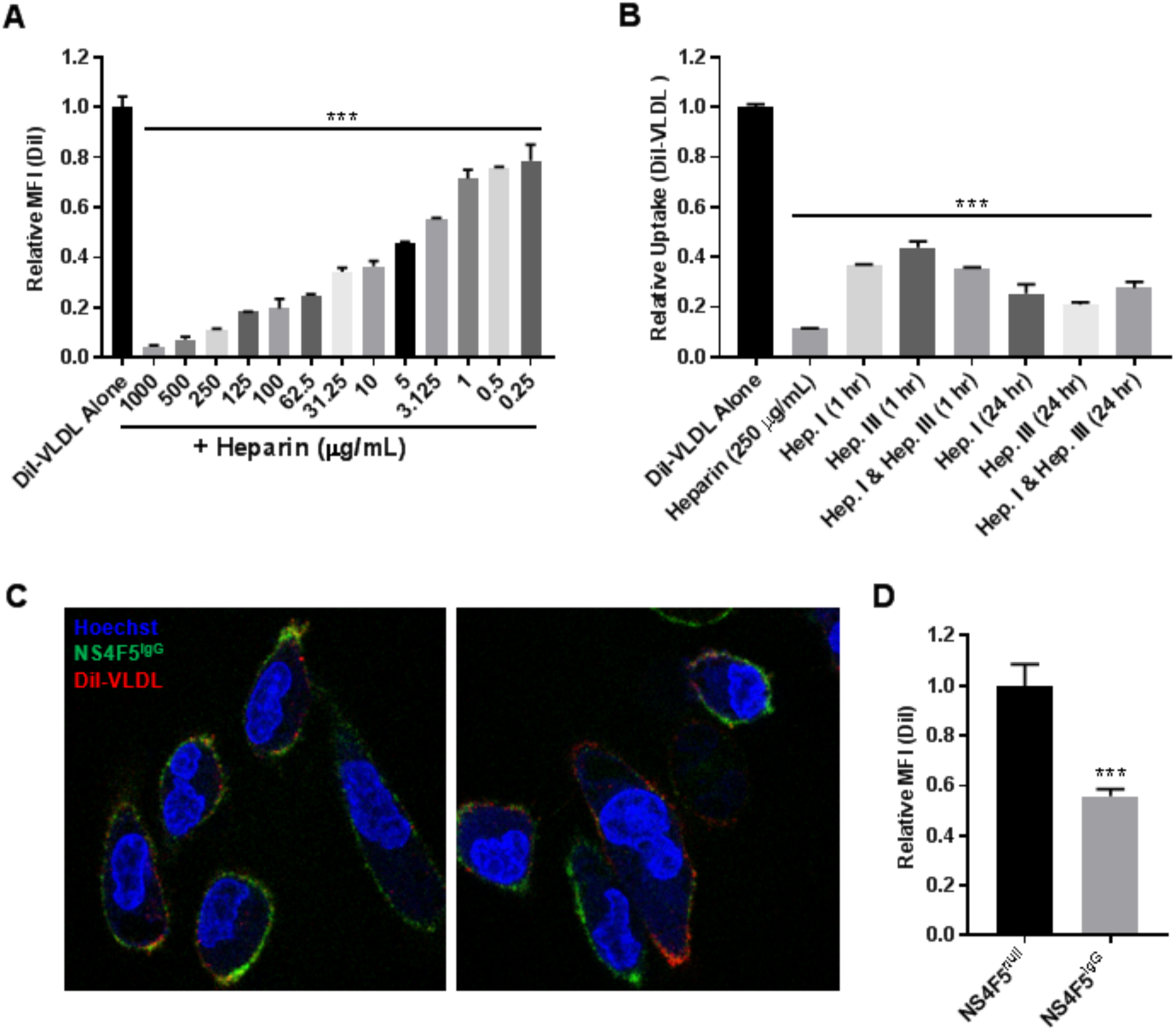
Lipoprotein uptake is reliant on HSPGs. DiI-VLDL binding and internalization is abrogated by treatment with heparin, heparinase, or an antibody directed against the HS^NS4F5^ motif in MDA-MB-231 BC cells. (**A**) Heparin treatment caused a significant, dose-dependent reduction in DiI-VLDL uptake, assessed by flow cytometry. One-way ANOVA with correction for multiple comparisons, ***p < 0.001. (**B**) Treatments with heparin or heparinase significantly reduced DiI-VLDL uptake as compared to the control, assessed via a plate-based DiI-VLDL uptake assay. One-way ANOVA with correction for multiple comparisons, ***p < 0.001, error as SD. (**C**) Representative confocal microscopy images of MDA-MB-231 co-labeled with Hoechst 33342 nuclear stain (blue), NS4F5 (green), DiI-VLDL (red). Both DiI-VLDLs and the HS^NS4F5^ antibody localize to the cell surface of BC cells incubated for 30 min at 4°C. (**D**) DiI-VLDL uptake was significantly reduced in MDA-MB-231 cells treated with unlabeled NS4F5^IgG^ antibody (3.2 µg/mL), as compared to those treated with the control NS4F5^null^ antibody. Two-tailed unpaired t-test with Welch’s correction, ***p < 0.001.

The effects of heparin and dynasore were tested across an array of BC cell lines (**Figure 4**). Despite differences in their characteristics (Table S1), all cell lines were shown to rapidly take up DiI-VLDLs. Binding and internalization were significantly reduced by heparin (500 µg/mL) or dynasore (80 µM), as quantified via flow cytometry and visualized using confocal microscopy (**Figure 4**).

**Figure 4.**
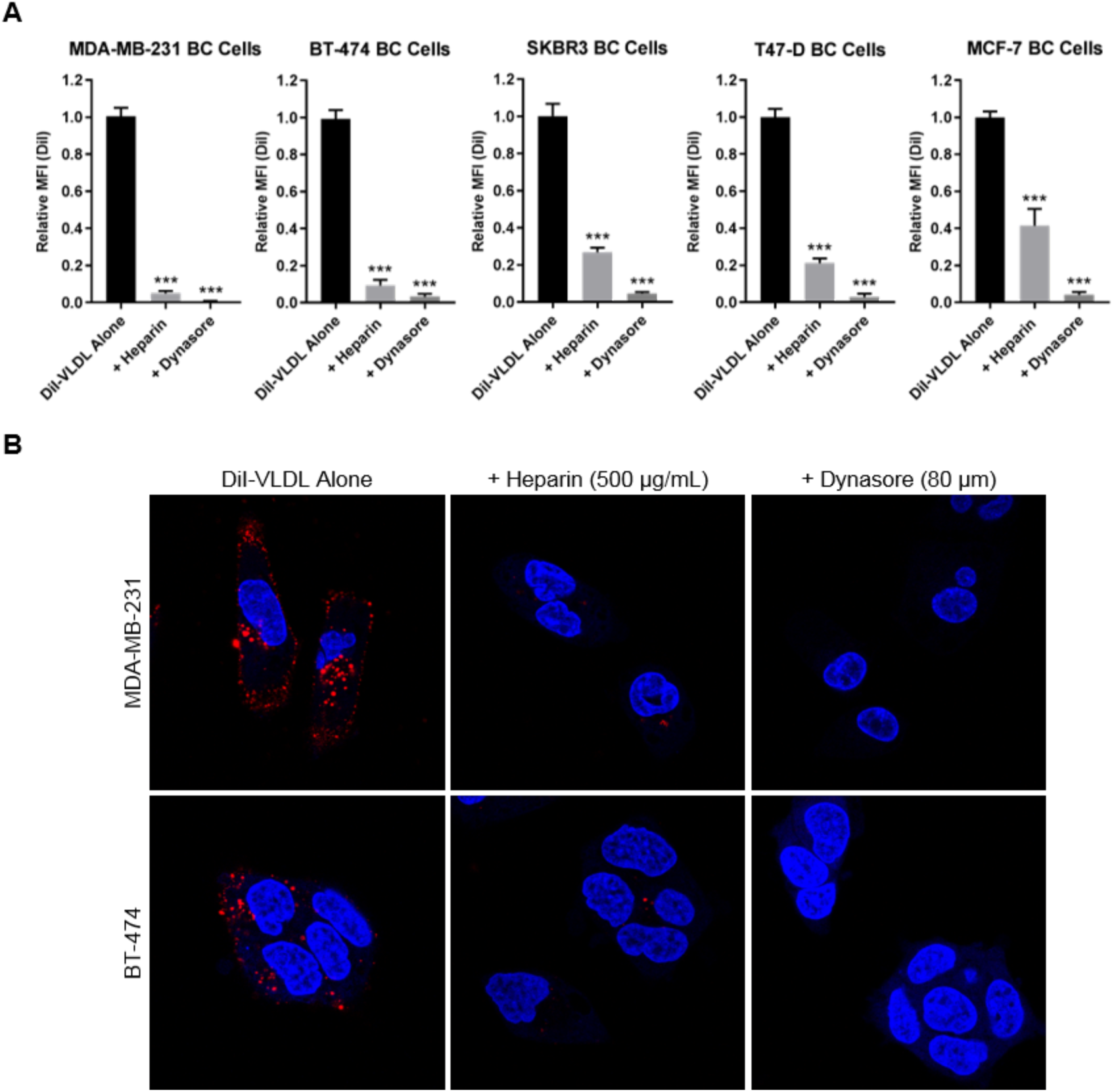
DiI-VLDL uptake is abrogated by treatment with heparin or dynasore across BC cell lines. (**A**) Treatment with heparin (500 µg/mL) or dynasore (80 µM) significantly reduced DiI-VLDL uptake, quantified by flow cytometry. Biological replicates from ≥ 3 experiments are shown. One-way ANOVA with correction for multiple comparisons, ***p < 0.001. (**B**) Effects of heparin (500 µg/mL) and dynasore (80 µM) on Di-VLDL binding and uptake in adherent BC cells visualized by confocal microscopy. Heparin or dynasore prevented both binding and internalization of DiI-VLDL particles at 37°C; DiI-VLDL (red), Hoechst nuclear stain (blue).

### A role for LPL and VLDLR in the uptake of VLDLs through receptor-mediated endocytosis

We used several genetic approaches to directly implicate LPL in the uptake of lipoprotein particles by BC cells. MCF-7 (LPL non-expressing) BC cells were engineered to overexpress LPL using a CMV promoter. MCF-7 pooled and clonal LPL-overexpressing cell lines displayed upregulated LPL mRNA expression (**Figure 5A**) and cell surface protein levels (**Figure 5B**) compared to pCMV Neo controls. Cell surface HS^NS4F5^ was significantly elevated in LPL-overexpressing MCF-7 cells, suggesting a direct relationship between expression of LPL and display of this LPL-binding motif (**Figure 5C**). Several other genes associated with lipid uptake, including CD36, LMF1 (lipase maturation factor 1), and VLDLR were also upregulated, with no differences observed in the expression of FA synthesis genes, (ACACA (acetyl-CoA carboxylase α), ACLY (ATP citrate lyase), FASN) or cholesterol synthesis genes (HMGCR (3-hydroxy-3-methylglutaryl-CoA reductase)) (**Figure 5D**, **E**). The lipid droplet (LD) content of LPL-overexpressing and Neo control MCF-7 cells was investigated both at baseline and with VLDL supplementation using LipidTox Neutral Red Lipid Stain. LPL-overexpressing cells displayed higher basal LD staining than pCMV Neo control cells, as shown by confocal microscopy using the MCF-7 pooled cell line pair (**Figure 5F**). VLDL supplementation (72 h, 100 µg/mL) increased the size and abundance of lipid droplets present in pCMV Neo and pCMV LPL cells, with the scope of change and LD content in LPL-overexpressing cells exceeding that of the Neo control (**Figure 5F**). This trend was consistent across media types, including matched FBS and LPD FBS-RPMI-1640, and was mirrored in the clonal cell lines (not shown). Together, these data indicate that overexpressing LPL alters the metabolism of MCF-7 BC cells by augmenting their capacity to take up lipids and ultimately increasing intracellular LD content.

**Figure 5.**
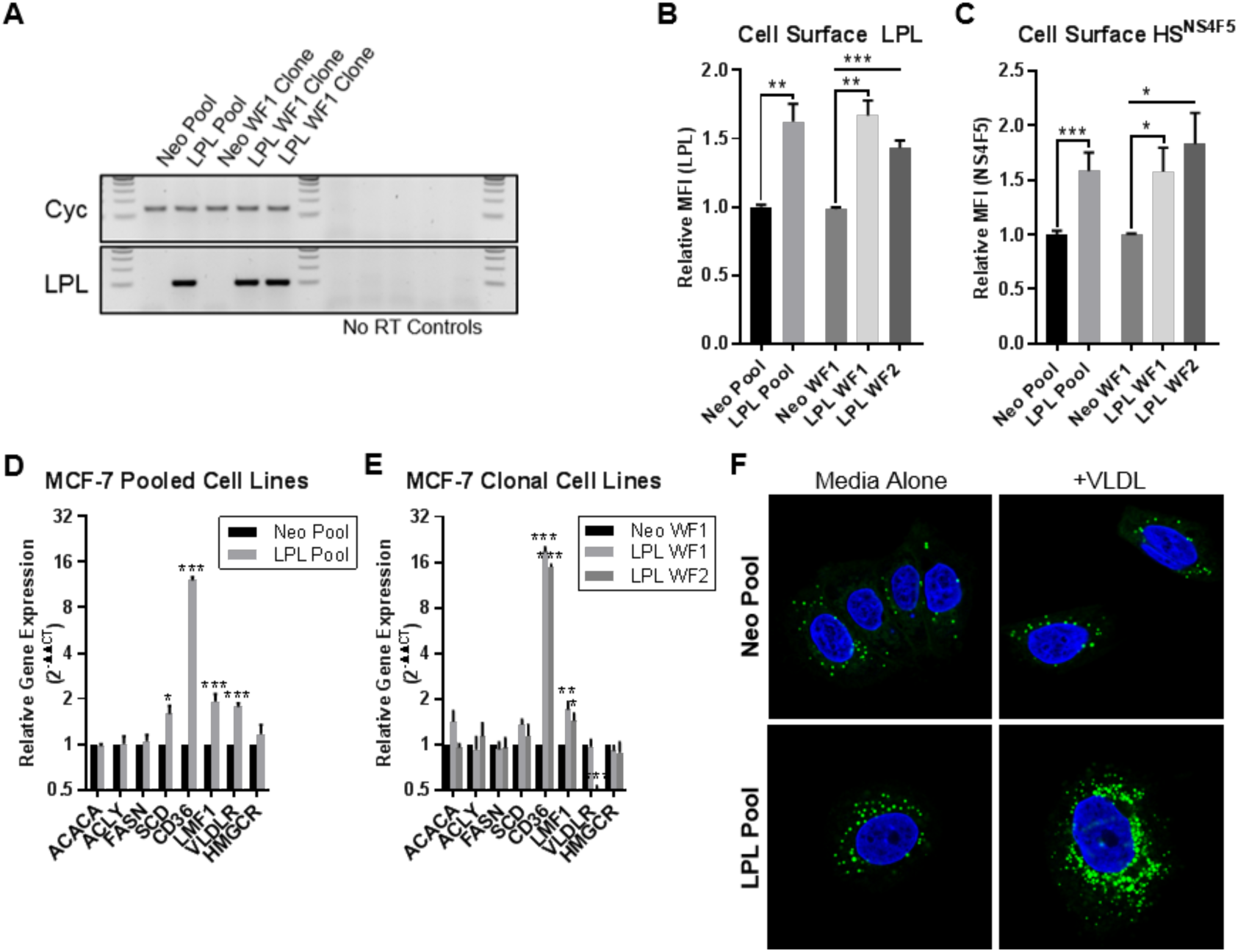
MCF-7 pCMV LPL cell lines overexpress LPL (RNA and protein), display increased cell surface HS^NS4F5^, and upregulate mRNA expression of genes involved in FA uptake. (**A**) RT-PCR products visualized on agarose gels. Increased LPL product was observed in MCF-7 pCMV LPL pooled and clonal cell lines compared to the pCMV Neo controls. (**B**) Increased cell surface staining of LPL in MCF-7 LPL-overexpressing cells quantified by flow cytometry. (**C**) MCF-7 LPL-overexpressing lines display increased cell surface HS^NS4F5^. (**D**-**E**) qRT-PCR of FA metabolism genes; relative gene expression calculated using 2^-ΔΔCT^ method. Data from ≥ 3 experiments; error is SEM. Statistical significance was determined using two-tailed unpaired t*-*tests. N.S. unless otherwise indicated; *P < 0.05, **p < 0.01, ***p < 0.001. (**F**) MCF-7 pCMV Neo and pCMV LPL cell pools were cultured for 72 h standard media +/- VLDL (100 µg/mL). MCF-7 LPL overexpressing cells exhibited higher LD content both at baseline and with VLDL supplementation. LipidTox (LD stain, green), Hoechst 3342 nuclear stain (blue).

We next investigated the effects of LPL knockdown by generating MDA-MB-231 LPL shRNA cell lines. As shown by qRT-PCR, LPL shRNA A cells exhibit a complete knockdown of LPL mRNA, whereas LPL shRNA D cells were partially knocked down, compared to the control (Figure S4A-B). Knockdown of LPL in the LPL shRNA A cells resulted in the transcriptional upregulation of FA synthesis genes, as well as an increase in the expression of CD36, VLDLR, LMF1, and HMGCR (Figure S4B). Partial knockdown of LPL mRNA (in LPL shRNA D cells) was associated with upregulation of the VLDLR mRNA alone. MDA-MB-231 control and LPL shRNA cells were compared for DiI-VLDL uptake and lipid droplet composition basally and with VLDL supplementation (100 µg/mL). To our surprise, LPL shRNA D cells (with partial knockdown of LPL mRNA, upregulation of VLDLR mRNA) displayed a significant increase in DiI-VLDL uptake (Figure S4C), as well as a higher lipid droplet content both basally and with VLDL supplementation, compared to LPL shRNA A cells or the Scr control (not shown).

Acute knockdown of LPL using siRNA was employed to avoid the compensatory changes in FA metabolism-related gene expression that occurred with stable LPL knockdown. We compared the effects of two siRNA constructs directed against each LPL and VLDLR in MDA-MB-231 cells to that of a scrambled control. As shown by qRT-PCR both LPL siRNAs efficiently reduced expression of LPL (**Figure 6A**). VLDLR siRNA constructs 1 and 2 significantly knocked down VLDLR gene expression compared to the control, and unexpectedly caused a concurrent reduction in LPL gene expression (a finding that was repeated in multiple experiments). All MDA-MB-231 siRNA-treated cells had significant reductions in DiI-VLDL uptake as compared to the control (**Figure 6B**). These effects were less dramatic in LPL siRNA 1 and 2 cells, where LPL alone was knocked down (and presumably present in the original tissue culture medium), and most significant in VLDLR siRNA 1 cells, which had the most efficient knockdown of both LPL and the VLDLR. While reduced expression of LPL was unexpected in the VLDLR siRNA-treated cells, these data support the hypothesis that *both* LPL and VLDLR are required for efficient lipoprotein uptake in MDA-MB-231 BC cells.

**Figure 6.**
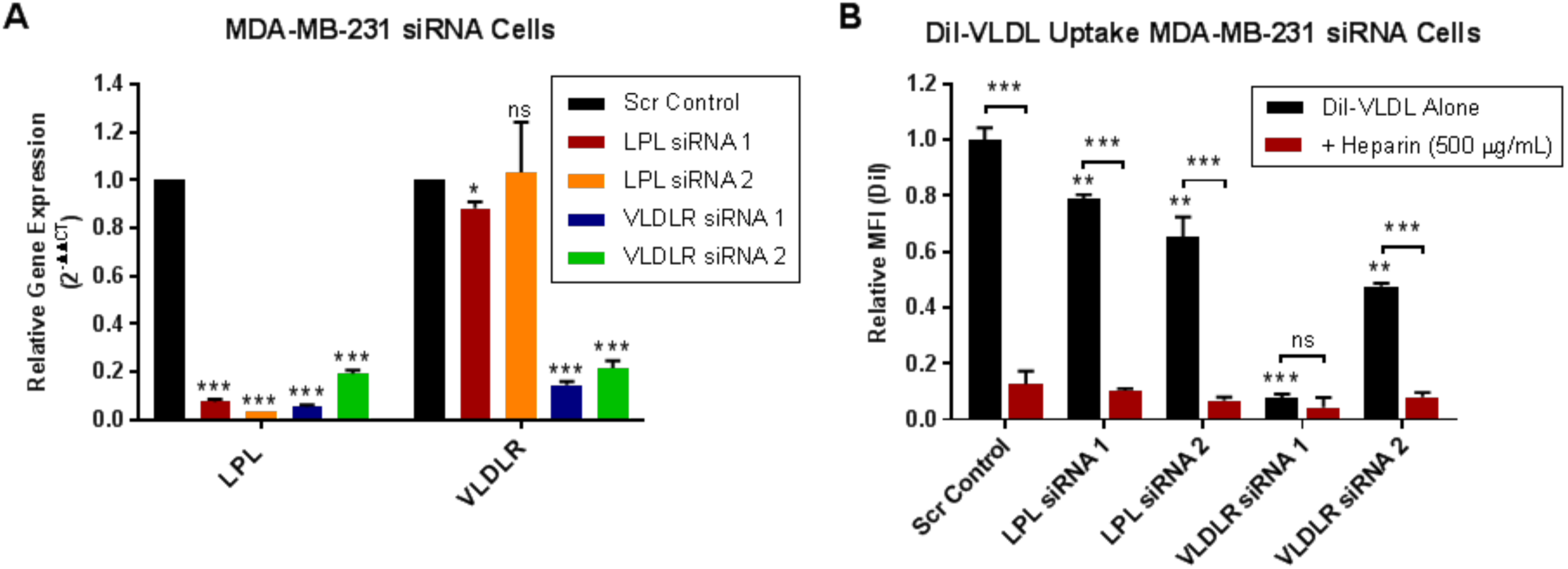
LPL and VLDLR are involved in VLDL uptake in MDA-MB-231 BC cells. (**A**) qRT-PCR verification of LPL and VLDLR knockdown in MDA-MB-231 BC cells treated with LPL or VLDLR siRNA. LPL and VLDLR gene expression of siRNA cells are relative to that of the Scr siRNA control. LPL siRNAs were knocked down for LPL mRNA, while VLDLR siRNAs significantly reduced both VLDLR and LPL mRNAs. (**B**) siRNA knockdown of LPL or VLDLR significantly reduces DiI-VLDL uptake in MDA-MB-231 BC cells. The greatest decreases were observed in VLDLR siRNA 1 cells, which had the highest efficiency knockdown of both VLDLR and LPL mRNAs. DiI-VLDL uptake was prevented by treatment with heparin in all cell lines except VLDLR siRNA, where the reduction from VLDL alone was not significant. P ≥ 0.05 (ns), *p < 0.05, **p < 0.01, ***p < 0.001.

### LPL supplementation increases DiI-VLDL uptake in BC cell lines

Our flow cytometry studies demonstrated that BC cells can take up exogenous LPL from sources such as the FBS of tissue culture media. We hypothesized that supplementing LPL at the time of DiI-VLDL uptake would increase lipoprotein binding and internalization by BC cells, and found that this was, in fact the case. Addition of bovine LPL (1 µg/mL) significantly increased DiI-VLDL uptake by MDA-MB-231 cells, and was unaffected by treatment with GSK264220A (GSK), an inhibitor of the catalytic activity of LPL [47, 48] (**Figure 7A**). This significant increase in DiI-VLDL uptake upon LPL supplementation was observed across BC cell lines (**Figure 7B**), albeit with variability potentially related to differences in basal LPL expression (BT-474 highest, MDA-MB-231; MCF-7 and T47-D as non-expressing cells). Visualization of MDA-MB-231 and BT-474 cells using fluorescence confocal microscopy revealed that addition of LPL increased both binding and internalization of VLDL particles (**Figure 7C**).

**Figure 7.**
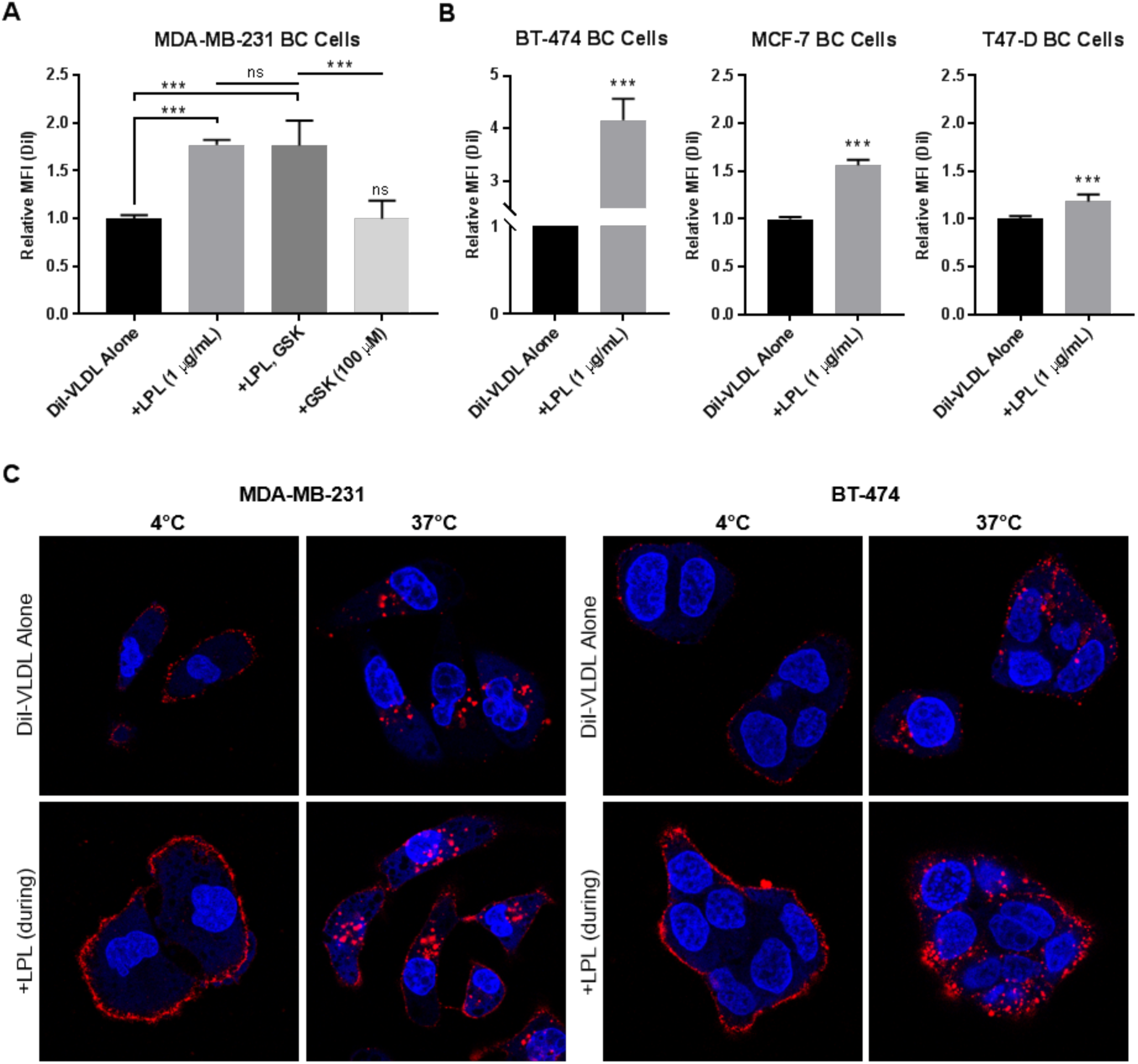
LPL supplementation increases DiI-VLDL binding and uptake in BC cells. (**A**) Quantification of DiI-VLDL uptake measured by flow cytometry. LPL (1 µg/mL) was supplied to MDA-MB-231 BC cells with DiI-VLDL (45 min, 5 µg/mL) at 37°C. LPL increased DiI-VLDL uptake (***p < 0.001, one-way ANOVA with correction for multiple comparisons). Treatment with lipase inhibitor GSK264220A did not significantly impact DiI-VLDL uptake. Biological replicates are displayed; error is SD. (**B**) LPL supplementation significantly increased VLDL uptake in LPL-expressing and non-expressing BC cell lines, as assessed by flow cytometry. Two-tailed unpaired t-test with Welch’s correction, *p < 0.05, **p < 0.01, ***p < 0.001. (**C**) LPL supplementation increases DiI-VLDL binding and internalization across BC cell lines. Representative confocal microscopy images of MDA-MB-231 and BT-474 BC cells incubated with DiI-VLDLs at 4°C or 37°C with or without LPL supplementation (1 µg/mL). DiI-VLDL (red), Hoechst 33342 nuclear stain (blue).

### Expression of FA metabolic genes in BC cells is impacted by the availability of lipoproteins in the media

MDA-MB-231 cells were cultured in RPMI-1640 media containing lipoprotein-depleted or matched control FBS for 96 h. qRT-PCR was then used to assess the expression of a panel of FA metabolism genes, including those involved with FA synthesis (ACACA, ACLY, ACSS2 (acyl-CoA synthase short chain family member 2), FASN, and SCD (stearoyl-CoA desaturase), lipoprotein/FA uptake (CD36, LPL, LMF1, VLDLR), cholesterol synthesis and uptake (HMGCR; LDLR), and lipid storage (DGAT1 (diacylglycerol acyltransferase-1), PLIN2 (perilipin 2)).

MDA-MB-231 BC cells cultured in LPDS media displayed significantly increased expression of genes involved in lipid synthesis (ACACA, ACLY, ACSS2, FASN, and SCD) and cholesterol metabolism (HMGCR and LDLR). Expression of FA/lipoprotein uptake genes, CD36 and LPL, was likewise increased following prolonged culture in LPDS media, while the expression of lipid storage genes, DGAT1 and PLIN2, was reduced (**Figure 8A**). While some cell line-specific changes were observed, these trends were largely consistent across a panel of BC cell lines (Figure S5). We selected three FA synthesis genes (ACLY, FASN, SCD) to address the question of whether VLDL addback to LPDS media could restore gene expression to the levels observed in the matched media control and found that this was the case. VLDL addback reduced the expression of all three FA synthesis genes back towards the level of the control baseline. MDA-MB-231 BC cells cultured in matched media supplemented with VLDL (100 μg/mL, 96 h) likewise exhibited decreased expression of ACLY, FASN, and SCD (**Figure 8B**).

**Figure 8.**
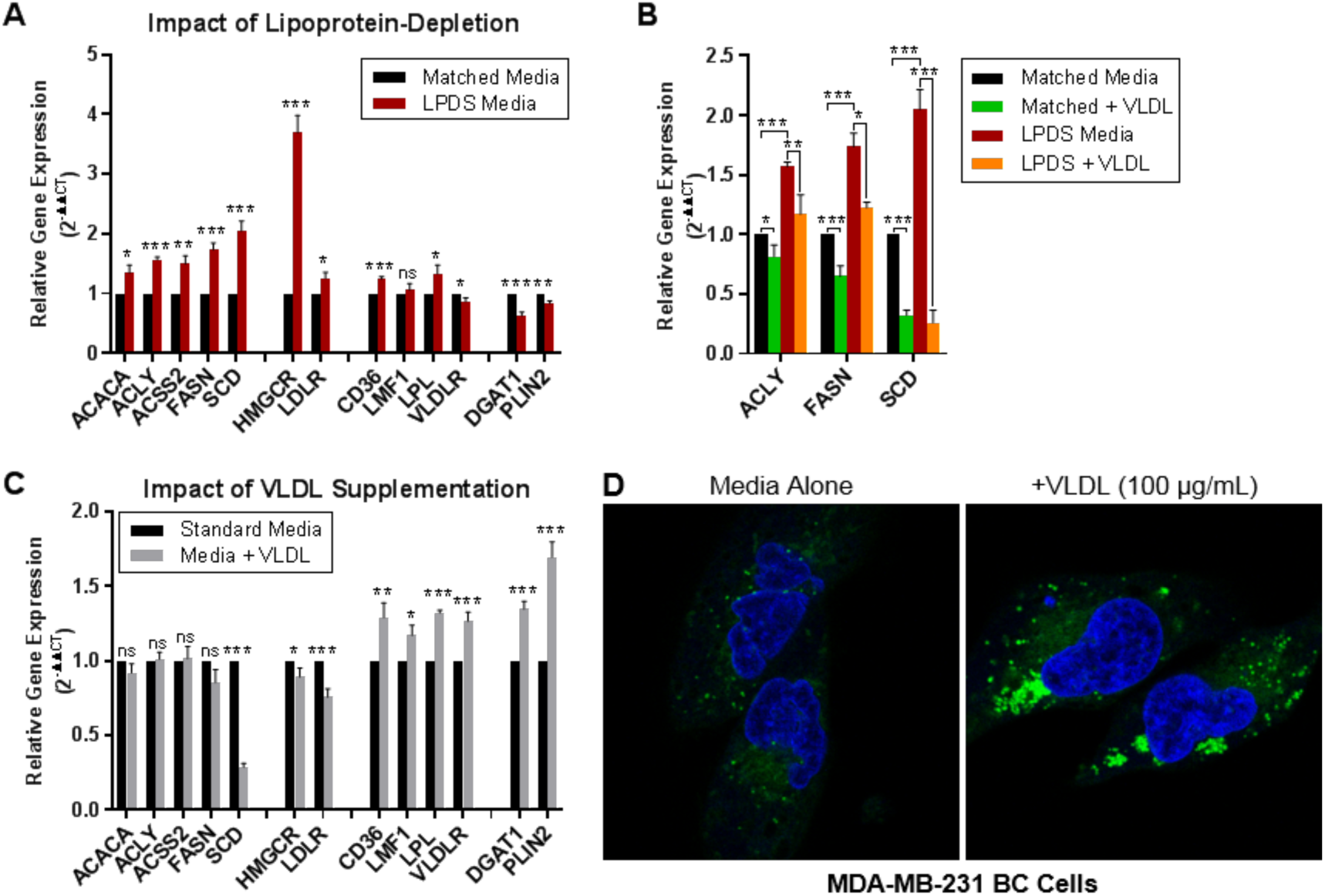
The presence of lipoproteins in the media impacts the expression of FA metabolic genes and abundance of lipid droplets in MDA-MB-231 BC cells. (**A**-**C**) qRT-PCR was used to assess the impact of media lipoproteins on the expression of FA metabolism genes. Relative gene expression is displayed as 2^-ΔΔCT^ (relative to the control). Data are mean +/- SEM of > 3 experiments. Two-tailed unpaired t*-*tests: P ≥ 0.05 (ns), *p < 0.05, **p < 0.01, ***p < 0.001. (**A**) MDA-MB-231 BC cells cultured in LPDS media for 96 h displayed significantly increased expression of FA synthesis and cholesterol metabolism genes, and significantly decreased expression of FA storage genes (DGAT1, PLIN2) compared to cells grown in matched control media. The expression of FA/lipoprotein uptake genes was variably impacted by lipoprotein depletion. (**B**) BC cells were grown for 96 h in matched or LPDS media supplemented or not with VLDLs (100 µg/mL) to assess the impact of supplementation or “addback” on the expression of select FA synthesis genes. Data are shown as relative gene expression normalized to the matched media control. Culturing cells in LPDS media resulted in a significant upregulation of ACLY, FASN and SCD (as shown in **A**). Addition of VLDLs (100 μg/mL) significantly reduced the expression of these genes back to or below the matched control media baseline. (**C**) qRT-PCR of cells grown in standard media supplemented or not with VLDLs (100 μg/mL, 72 h). VLDL supplementation was associated with increased expression of FA/lipoprotein uptake and FA storage genes; ns change or decreases in FA synthesis and cholesterol metabolism genes. (**D**) Increased size and abundance of intracellular LDs observed in cells grown in media supplemented with VLDLs (100 μg/mL, 72 h) compared to the media alone control. Cells were stained with the Hoechst 33342 nuclear stain (blue) and LipidTOX Red Neutral Lipid Stain (green) and visualized using fluorescence confocal microscopy.

We next explored the impact of VLDL supplementation (100 μg/mL, 96 h) on the gene expression profile of MDA-MB-231 BC cells cultured in standard RPMI-1640 media. Here, addition of VLDLs to the media was associated with increased expression of genes involved in lipid uptake (CD36, LMF1, LPL, VLDLR) and storage (DGAT1, PLIN2), significant reductions in reductions in the expression of SCD, HMGCR, and LDLR, and no significant changes in the expression of FA synthesis genes (ACACA, ACLY, ACSS2, FASN) (**Figure 8C**). We hypothesized that transcriptional upregulation of lipid uptake and storage genes may be reflected in the abundance of lipid droplets. Visualization of lipid droplets via confocal microscopy revealed that MDA-MB-231 BC cells cultured in media supplemented with VLDLs (100 μg/mL, 72 h) displayed a stark increase in the size and abundance of LDs compared to that of cells grown in media alone (**Figure 8D**). This was recapitulated in multiple cell lines (BT-474, MCF-7, T47-D; not shown). These data indicate that BC cells can respond to exogenous lipid availability and shift their metabolism-related gene expression programs (lipogenesis vs. uptake) to meet their FA requirements.

## DISCUSSION

We previously showed that a majority of breast tumors display both LPL and CD36, and that provision of VLDL particles to BC cells (in the presence of LPL) stimulates cell growth and enables cells to evade the cytotoxic effects of FA synthesis inhibition [3]. These results, in concert with the observation that exogenous FFA can rescue cancer cells from the cytotoxic effects of FA synthesis inhibitors [5, 49-53], prompted the idea that BC cells might deploy LPL in a manner analogous to that of nonmalignant LPL-producing cells, such adipocytes or cardiomyocytes. Our findings support a model wherein LPL is bound to the cell surface, both in BC cell lines and in clinical BC tissue. Several lines of evidence support the identity of the binding site as a HSPG molecule, including LPL-displaceability by heparin and significant reductions in DiI-VLDL binding and internalization following heparin or heparinase treatment. Decreased VLDL uptake by BC cells in the presence of an antibody directed against HS^NS4F5^ implicates this specific HSPG motif as the binding moiety. Notably, the HS^NS4F5^ sequence (GlcNS6S-IdoA2S)_3_ is repeated in heparin and was previously identified as an LPL-binding sequence [33] found on the surface of MCF-7 BC cells [54]. Though MCF-7 and several other BC cell lines do not express detectable levels of LPL mRNA, they do display HS^NS4F5^ on their cell surface. Our data indicate that this motif acts as a “bait” to acquire exogenous LPL, and that this LPL can facilitate the uptake of lipoproteins. Acquisition of LPL in this manner could be important in anatomic areas where primary or metastatic BC reside, such as the mammary fat pad or fatty bone marrow.

Our data do not exclude a hydrolytic role for LPL on the BC cell surface. Indeed, the cell lines that we have examined express CD36 and thus are equipped to benefit from extracellular FFA. On the other hand, our experimental findings clearly establish a noncatalytic role for the cell surface LPL-HS^NS4F5^ complex in the binding and internalization of VLDLs. The rapid and temperature-sensitive uptake of intact lipoproteins, and its prevention by dynasore support receptor-mediated endocytosis as the mechanism observed in our studies.

Many attempts were undertaken to implicate LPL in the endocytosis of VLDLs, in addition to the indirect evidence provided by experiments with heparin, heparinase, or anti-HS^NS4F5^. We learned through flow cytometry studies that cell lines that do not express LPL can acquire it from exogenous sources, such as the FBS in the media. Attempts to remove LPL from the system proved challenging, as removing it from media incited shifts in the metabolic phenotype of cells. While some variations were observed among BC cell lines, removal of lipoproteins generally induced the expression of FA and cholesterol synthesis-related genes, mirroring the metabolic plasticity observed by Daniels and coworkers in prostate cancer cells [52].

We generated LPL-overexpressing and shRNA knockdown cell lines in order to isolate a specific LPL phenotype. Enforced LPL expression in otherwise non-expressing MCF-7 cells caused prominent increases in the expression of CD36 and LMF1 mRNAs, as well as the abundance of HS^NS4F5^ on the cell surface, an apparent FA uptake program to capitalize on the availability of endogenously produced LPL. These changes were associated with increases in the size and abundance of LDs in these cells basally and upon supplementation with VLDL. On the other hand, shRNA knockdown of LPL in MDA-MB-231 BC cells induced the expression of mRNAs coding lipogenic enzymes (ACACA, ACLY, FASN, SCD), as well as VLDLR and CD36. We interpret this as a shift towards FA synthesis and compensatory upregulation of other genes in the lipid uptake program. Acute knockdown of LPL by siRNA caused a modest reduction of VLDL uptake, owing presumably to the presence of residual LPL from the FBS of the media. Ultimately, the clearest evidence of LPL involvement came from a series of simple experiments showing that LPL supplementation at the time of DiI-VLDL treatment causes significant increases in DiI-VLDL binding and internalization. This trend was consistent across all cell lines tested, directly implicating LPL, and further validating the ability of cancer cells to take up and rapidly use exogenous LPL to facilitate lipoprotein uptake.

While it was feasible that HSPG-bound LPL performed this non-catalytic function alone, our data provide evidence that the VLDLR is also involved. Like LPL, the VLDLR is expressed to a variably by the cancer cell lines used in these studies; its expression appears to be impacted by exogenous lipid supply, as well as the status of LPL transcriptional machinery. siRNA knockdown of the VLDLR caused a concurrent reduction in LPL gene expression (a surprising finding that was repeated with different siRNAs over multiple experiments), and in the most efficacious dual-knockdown condition, nearly eliminated VLDL uptake. While the exact nature of the VLDLR-LPL interaction warrants future study, associations between LPL and the VLDLR are well established in other contexts.

VLDLR is highly expressed on the capillary endothelium of skeletal muscle, heart, adipose and brain tissue, and by macrophages, but only to a limited extent in liver [55, 56]. VLDLR binds to apolipoprotein E (apoE)-containing lipoproteins, including VLDL [57-59]. Studies of the distribution and ligand specificity of VLDLR revealed marked similarities with LPL, prompting speculation that the VLDLR worked in concert with LPL to facilitate the binding of TG-rich lipoproteins and delivery of FFAs to tissues [59-61]. This hypothesis has been supported by almost two decades of research. There are now at least three non-mutually exclusive configurations in which LPL and VLDLR can interact: (1) LPL can bind directly to the VLDLR, increasing the binding affinity and subsequent internalization of TG-rich lipoproteins through receptor-mediated endocytosis, (2) HSPG-bound LPL can facilitate interactions between lipoprotein and the VLDLR, analogous to cell surface binding of heparin-binding growth factors and their receptors, and (3) LPL mediated-hydrolysis of VLDLs can lead to the formation of smaller (ApoE-rich) remnant particles, which may remain bound to LPL before being recognized and endocytosed by receptors [62]. The high-capacity model for lipoprotein uptake in BC cells outlined herein involves HSPG-bound cell surface LPL binding to VLDLs and facilitating the uptake of intact lipoproteins via receptor-mediated endocytosis in concert with the VLDLR (**Figure 9**).

**Figure 9.**
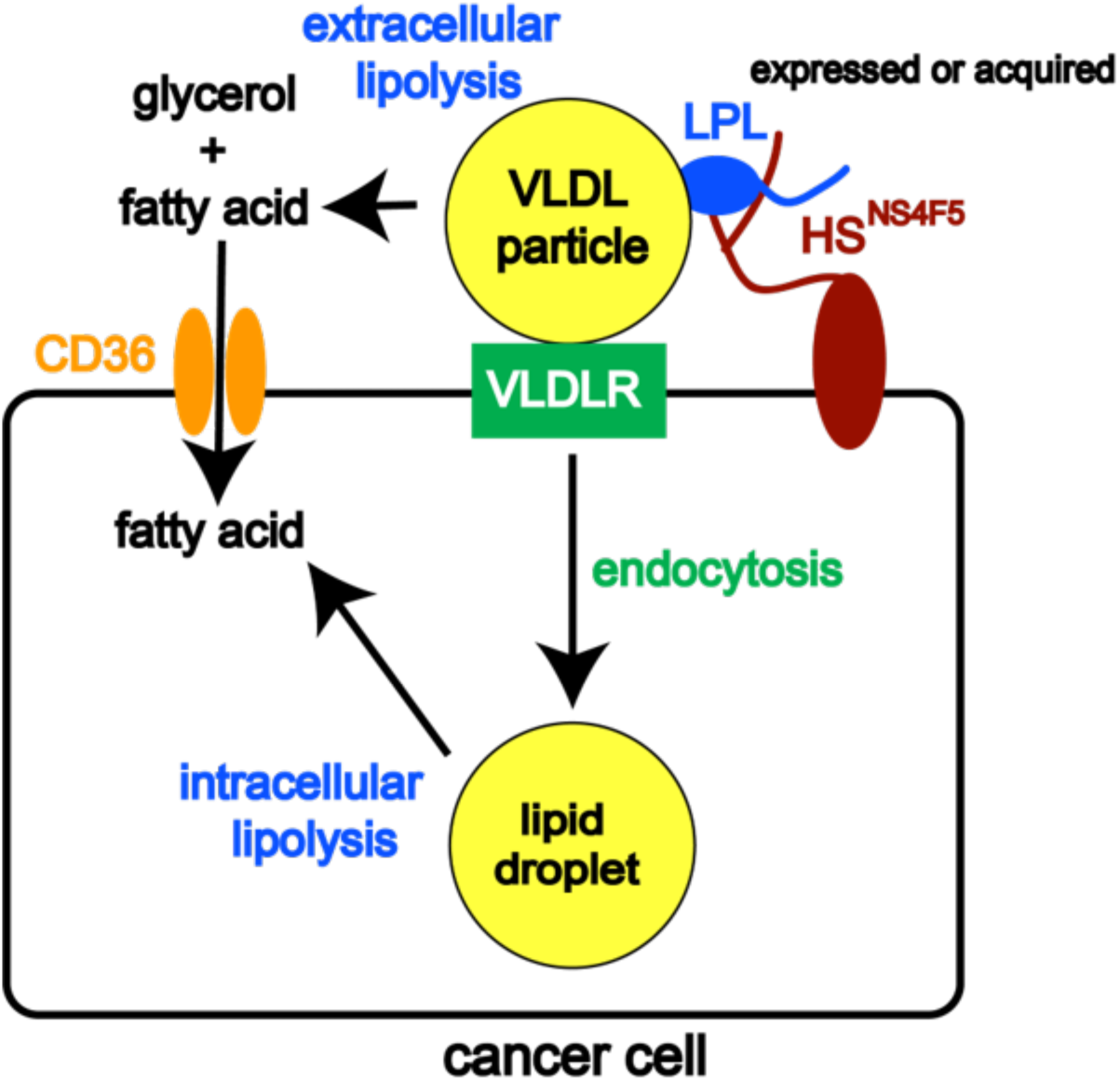
Model of lipoprotein uptake in BC cells. Our previous and current work has shown that LPL binds to a specific HSPG motif (HS^NS4F5^) on the cancer cell surface, where it can function in two distinct roles: (1) catalyzing the hydrolysis of TG-rich lipoproteins, creating FFA that can enter the cell through FA uptake channel, CD36, or (2) acting non-catalytically in concert with VLDLR to facilitate the uptake of intact VLDL particles by receptor-mediated endocytosis. LPL can be synthesized or acquired from exogenous sources, such as media FBS *in vitro* or LPL-secreting tissues (e.g. adipocytes or macrophages) *in vivo.* By functioning in this dual-capacity, LPL plays a prominent role in mediating the uptake of the lipids and FFAs required to sustain the growth and proliferation of cancer cells.

We recently pointed out that the literature related to lipid metabolism and cancer has emphasized the role of FA synthesis and proposed that uptake of exogenous lipid is also important [2]. Here, we have elucidated a novel mechanism through which BC cells take up VLDL particles. Our data support the involvement of LPL, the VLDLR, and HSPGs in the endocytosis of intact lipoproteins, and leave the potential for LPL to also act enzymatically at the cell surface. We add to the growing body of evidence supporting the importance of exogenous lipid uptake in cancer cells, broadly [63, 64], and LPL-mediated uptake of lipids in cancer, more particularly. While investigations into the role of LPL have been most prominent in CLL [12, 17-19, 21, 23, 24, 65], LPL has emerged as a protein of interest in a variety of different tumor types, including breast cancer [3, 66, 67], cervical squamous cell carcinoma [68], hepatocellular carcinoma [5], gastric cancer [69-71], colorectal cancer [9] lung cancer [6, 10, 72, 73], and prostate cancer [74].

Our findings have implications for the development of cancer therapeutics aimed at lipid metabolism. We and others have shown that cancer cells display increased sensitivity to FA synthesis inhibitors when cultured in lipid-reduced or lipoprotein-depleted media [5, 51-53]. It has likewise been demonstrated that cancer cells may adjust their relative reliance on FA synthesis vs. lipolysis/uptake in response to external factors, such as nutrient availability [52]. Our data add to the mounting evidence that targeting FA synthesis may not achieve maximal efficacy in certain tumors, including BC, because the they may employ both FA synthesis and uptake, and the reliance on these pathways may shift in response to altered nutrient availability or therapeutic inactivation of only one of them.

## EXPERIMENTAL PROCEDURES

### Cell lines and tissue culture

MCF-7, MDA-MB-231, BT-474, DU4475, SKBR3 and T47-D BC cells, and HeLa cervical cancer cells were acquired from the American Type Culture Collection (ATCC) and cultured in phenol red-containing HyClone RPMI-1640 media with 10% (v/v)-heat-inactivated FBS (GE Healthcare Life Sciences) and 1% penicillin-streptomycin. MCF10A mammary epithelial cells were cultured in DMEM/F12 growth media (Invitrogen) supplemented with 5% horse serum (Invitrogen), 20 ng/mL epidermal growth factor (Peprotech), 0.5 mg/mL hydrocortisone, 100 ng/mL cholera toxin, 10 µg/mL insulin (Sigma), and 1% penicillin-streptomycin. All cells were maintained at 37°C in a humidified atmosphere containing 5% CO_2_. Characteristics of the cell lines used are in Table S1. Lipoprotein-depleted (LPD) and “matched control” FBS were obtained from Kalen Biomedical (Cat. No. 880100; 880170).

### Manipulated cell lines

#### MDA-MB-231 LPL shRNA cells

MDA-MB-231 shRNA cells were generated through lentiviral transduction using HuSH shRNA plasmid panels (29 mer) with pGFP-C-Lenti vectors and the Lenti-vpack Packaging Kit (TR30037) according to manufacturer guidelines (OriGene Technologies, Rockville, MD). shRNA plasmids included four LPL shRNAs and a negative control (TL311692). Cells stably transduced with the shLPL or scrambled shRNA construct were selected in puromycin. Successful transduction and selection were verified through visualization of the GFP tag and qRT-PCR.

#### MDA-MB-231 siRNA cells

siRNA constructs targeting LPL or VLDLR (Sigma) were transfected into MDA-MB-231 BC cells using Lipofectamine RNAiMAX (ThermoFisher Scientific) according to manufacturer guidelines. The following plasmids were used: LPL siRNA 1 (SASI_Hs01_00208454), LPL siRNA 2 (SASI_Hs01_00208455); VLDLR siRNA 1 (SASI_Hs02_00335553), VLDLR siRNA 2 (SASI_Hs01_00219062); MISSION siRNA Universal Negative Control #1, SIC001 (Sigma-Aldrich). Samples were assayed for DiI-VLDL uptake and RNA was collected for RT-PCR validation of the knockdown on day 4 of the experiment.

#### MCF-7 pCMV -Neo and -LPL cells

Overexpression of LPL was performed using a human LPL clone in pCMV-6AC plasmid vector synthesized by OriGene (Rockville, MD; Cat. No. SC322258). An empty pCMV-6AC (“pCMV Neo”) was included as the control. Plasmids were transfected using Lipofectamine 2000 DNA transfection reagent (ThermoFisher Scientific). Pooled and clonal cell lines were selected by limiting dilution in neomycin (G418) and propagated in G418-containing media. Total RNA was subjected to on-column DNase digestion with the RNase-free DNase set (Qiagen, Cat. No. 79254), cDNA synthesis and RT-PCR, as previously described.

### Materials

The following reagents were prepared as stock solutions in DMSO and stored at −20°C until use: GSK264220A, lipase inhibitor (50 mM; Cayman Chemical, Ann Arbor, Michigan); Dynasore, inhibitor of dynamin 1 and 2 (80 mM; Sigma-Aldrich); Hoechst 33342 nuclear stain (20 mM; ThermoFisher Scientific). The remaining reagents were prepared and stored at 4°C until use: heparin sodium salt from porcine intestinal mucosa (50 mM stock solution in H_2_O; Sigma, Cat. No. H3149), human DiI-VLDLs (1 mg protein/mL; Alfa Aesar Chemicals), LPL from bovine milk (Sigma-Aldrich, Cat. No. L2254). The protocol for preparing LPL from suspension (3.8 M ammonium sulfate, 0.02 M Tris HCl, pH 8.0) was adapted from Nishitsuji, Hosono [75]. Heparinase I from *Flavobacterium* (Sigma) was dissolved at 1 mg/mL in 20 mM Tris-HCl, pH 7.5, 50 mM NaCl, 4 mM CaCl_2_, and 0.01% BSA. Heparinase III from *Flavobacterium* (Sigma) was reconstituted in 20 mM Tris-HCl, pH 7.5, 4 mM CaCl_2_, and 0.01% BSA.

### Antibodies

Our mouse monoclonal anti-human LPL antibody described in [3] was conjugated to Alexa-Fluor 647 (AF-647) using the AF-647 antibody labeling kit (ThermoFisher Scientific, Cat No. A20186). A monoclonal antibody directed against the HS^NS4F5^ HSPG motif, (GlcNS6S-IdoA2S)3 was supplied by Nicole Smits. The binding site from the single chain NS4F5 antibody (NS4F5^scFv^) [45] was converted to a functional full-length human IgG2 antibody (NS4F5^IgG^). Briefly, the VH region of NS4F5^scFv^ was linked to the human γ-2 constant region under the control of the human T lymphotropic virus elongation factor-1 promoter, whereas the VL region of NS4F5^scFv^ was linked to the C-gene of the human lambda constant region under control of a cytomegalovirus promoter. A non-functional isotype control (NS4F5^null^) was generated by modifying the VH sequence of NS4F5^IgG^ (CARSGRKGRMR to CARSGSGGSGS). Heavy and light chain constructs of NS4F5 (NS4F5^IgG^, NS4F5^null^) were co-transfected into HEK-293 cells. Antibodies were purified by protein A-sepharose affinity chromatography, buffer exchanged into PBS, and analyzed using SDS-Page and size exclusion chromatography HPLC analysis using a TOSOH TSK gel SuperSW3000 column. Endotoxin levels were < 2 EU/mg.

### Quantitative RT-PCR

Total RNA was isolated using RNeasy minicolumns from cell extracts prepared with QiaShredder (Qiagen RNeasy Kit). The concentration and purity of RNA were assessed by a NanoDrop DM-1000 spectrophotometer (NanoDrop Technologies). cDNA was produced using iScript Reverse Transcription Supermix for RT-qPCR (BioRad) with 1 μg RNA template. Quantitative RT-PCR was performed using TaqMan Gene Expression Master Mix and TaqMan Gene Expression Assays (ThermoFisher Scientific). Information for primers is in Supplemental Table S2. qRT-PCR was performed on a Bio-Rad CFX 96 Real Time System C1000 Thermal Cycler with the following program: (1) 50°C× 2 min, (2) 90°C× 10 min, (3) 95°C x 15 sec, (4) 60°C × 1 min + plate read, (5) Go to (3) 39 more cycles, (6) Melt curve 65°C to 95°C increment 0.5°C for 5 sec + plate read. Relative gene expression results are displayed in accordance with the 2^-ΔΔCT^ method (Livak and Schmittgen 2001), with the expression of target genes first normalized to the housekeeping gene cyclophilin. Where applicable, RT-PCR products were separated on 2% agarose gels prepared using 0.5x TBE with SYBR Safe DNA gel stain (Invitrogen) and visualized using a Bio-Rad Molecular Imager ChemiDoc XRS+ with Image Lab Software.

### Flow Cytometry

Cells were detached with CellStripper non-enzymatic cell dissociation reagent (Fisher Scientific). Viability was assessed using the Molecular Probes LIVE/DEAD stain kit with excitation at 405 nm (20 min, 1:3000 dilution). Cells were blocked with human IgG (1.25 mg/mL, 15 min; Sigma Aldrich). Surface staining of unfixed/unpermeabilized cells employed the following antibodies: mouse monoclonal LPL-AF-647 (1 µg/mL, 30 min); Dylight 650-NS4F5^IgG^ or Dylight 650-NS4F5^null^ (3.2 µg/mL, 30 min). All incubations were carried out at 4°C with PBS washes between each step. Analysis of intracellular antigens was achieved using the Foxp3 / Transcription Factor Staining Buffer Set (ThermoFisher Scientific). Following staining, cells were fixed with 0.5% formalin (20 min). Fluorescence was measured using a FACScan system on a MacsQuant-10 Flow Cytometer (Miltenyi Biotec) and data were analyzed using FlowJo 10 software (FlowJo, LLC). For LPL staining, the unstained control condition was verified as having equivalent fluorescent reads as a mouse IgG2b K Isotype Control APC antibody (ThermoFisher Scientific).

### VLDL Uptake Experiments

#### DiI-VLDLs

VLDLs labeled with DiI (1,1’-dioctadecyl-3,3,3’-tetramethyl-indocarbocyanine perchlorate) were purchased from Alfa Aesar (Cat. No. J65568) and stored at 4°C until use. VLDLs with protein- and lipid-specific labels were prepared using a technique adapted from Goulbourne et al. [37]. Briefly, DiI-VLDLs were dialyzed into PBS containing 0.05 M sodium borate (pH 8.5) and incubated with DyLight 650 NHS-Ester (ThermoFisher Scientific) at room temperature for 1.5 h. Non-reacted DyLight dye was removed by extensive dialysis against PBS.

#### DiI-VLDL plate-based uptake assay

MDA-MB-231 cells were seeded subconfluently in 12-well plates and allowed to adhere. Growth media were replaced with RPMI-1640 containing 10% LPDS (Kalen) overnight, and cells were assayed for uptake of DiI-VLDL using methods adapted from Teupser et al. [76]. Briefly, cells were pre-treated for 30 minutes in SFM prior to a 1 h incubation with 10 µg/mL DiI-VLDL. Surface-bound DiI-VLDL were removed by twice treating with acid wash buffer (0.5 M acetic acid with 150 mM NaCl, pH 2.5). Cells were then washed with DPBS with calcium and magnesium, lysed in 1% SDS, 0.1 M NaOH, transferred to a black 96-well half-area plate (Greiner Bio-One) and assessed using a SpectraMax i3x microplate reader (Molecular Devices; Ex/Em 520/580 nm). Fluorescence was corrected for protein content as measured by BCA assay (Thermo Scientific).

#### DiI-VLDL flow cytometry-based uptake assay

Cells were plated in complete media into 12-well plates, allowed to adhere overnight, and serum starved for 1-8 h. Confluency was less than 70% at the time of treatment. All pre-treatments (30 min) and VLDL uptake incubations (45 min) were carried out at 37°C in a humidified atmosphere containing 5% CO_2_, unless otherwise indicated. Cells were washed extensively and then harvested by extended trypsinization. Samples were run on a ZE-5 flow cytometer (Bio-Rad). Data were analyzed using FlowJo 10 software (FlowJo, LLC).

### Confocal Microscopy

#### Lipoprotein uptake

Cells were seeded subconfluently into an 8-chambered Lab-Tek coverglass (Nunc), allowed to adhere overnight, serum starved for 1-8 h, and then assessed for DiI-VLDL uptake. Treatments with uptake-inhibiting reagents (30 min) were followed by a 45 min incubation with DiI-VLDL (5 µg/mL). Cells were rinsed with DPBS with calcium and magnesium, fixed with 3% paraformaldehyde in DPBS. Nuclei were stained with DAPI (Thermo Scientific) or Hoechst 33342 (Sigma-Aldrich). Images were acquired by confocal fluorescence microscopy with a Zeiss LSM 800 microscope equipped with a 63X oil-immersion objective with identical exposure settings for all experimental conditions. Images were processed uniformly across comparisons with Zen Lite 2.3 (Zeiss) and ImageJ 1.49 (Fiji).

#### Lipid droplet staining

Cells were grown in chambered Lab-Tek coverglass wells (Nunc), washed with DPBS, stained with Hoechst 33342 (1 µg/mL), fixed in 3% paraformaldehyde in DPBS with calcium and magnesium, and then incubated with LipidTOX Red Neutral Lipid Stain (Invitrogen; 1:800, 30 min at 37°C).

### Lipase Assay

Lipase activity was measured using a fluorescent assay [77] using Zwitterionic detergent 3-(N,N-Dimethylmyristylammonio) propanesulfonate and EnzChek Lipase Substrate, green fluorescent, 505/515 (Sigma-Aldrich; 250 µM stock). Fluorescence was measured on a SpectraMax i3x plate reader and SoftMax Pro software from Molecular Devices (Ex/Em: 482 nm/515 nm, 495 nm filter cutoff). Assays were carried out after 10 min incubation at 37°C in black half area 96-well plates (Greiner Bio-One) in 0.15 M NaCl, 20 mM Tris-HCl, pH 8.0, 0.0125% Zwittergent, and 1.5% FA-free BSA in a total volume of 100 µL. The average of no lipase controls was subtracted from raw values to account for background hydrolysis.

### Cell Viability Assays

Cells were seeded into 96-well plates, allowed to adhere overnight, treated with drug formulations, and cultured for 72 h at 37°C before viability assay using the ATP-based CellTiter-Glo (CTG) 2.0 Assay (Promega). Luminescence was read with the LMAX II luminometer (Molecular Devices). Background luminescence was calculated from CTG alone wells. Luminescence values (RLU) were normalized to vehicle and compiled as % DMSO control. IC_50_ determination was performed using a non-linear regression curve-fitting algorithm: log(inhibitor) vs. response-variable slope (four parameter) with Graph Pad Prism 6.0.

### Cell death and apoptosis assays

Apoptosis was assessed using the Dead Cell Apoptosis Kit with Annexin V Alexa Fluor 488 and Propidium Iodide (PI) (ThermoFisher Scientific). Flow cytometry data obtained with the 8-color MACSQuant-10 (Miltenyi Biotec) were analyzed using FlowJo software. Percent apoptosis was the percentage Annexin V+ cells in the population.

### Statistics

Statistical significance was evaluated using either an unpaired two-tailed student’s t-test with Welch’s correction or one-way ANOVA with correction for multiple comparisons (Dunnett’s multiple comparisons test), where applicable. P-values < 0.05 were deemed significant: *p<0.05, **p<0.01, ***p<0.001. Error presented as mean ± SD or SEM, as indicated.

## Acknowledgements

We thank Cary N. Mariash for his critical reading of this manuscript.

## FOOTNOTES

This work was supported by NIH grant RO1CA58961 (to WBK) and a Norris Cotton Cancer Center grant (to WBK); C16/15/073 and C32/17/052 from the KU Leuven (to JVS) and Interreg V-A EMR23 EURLIPIDS (to JVS). The authors acknowledge the following Shared Resources Facilities: Immune Monitoring and Flow Cytometry Resource (IMFCSR), Irradiation, Pre-clinical Imaging and Microscopy Resource (IPIMSR) at the Norris Cotton Cancer Center at Dartmouth with NCI Cancer Center Support Grant 5P30CA023108-37, and support from the National Institute of General Medical Sciences COBRE Grant (P30GM103415-15).

## Abbreviations Employed

ACACA: acetyl-CoA carboxylase α
ACLY: ATP citrate lyase
ACSS2: acyl-CoA synthase short chain family member 2
BC: breast cancer
CD36: fatty acid translocase
CLL: chronic lymphocytic leukemia
DGAT1: diacylglycerol acyltransferase-1
FA: fatty acid
FASN: fatty acid synthase
FFA: free fatty acid
GPIHBP1: glycosylphosphatidylinositol-anchored high-density lipoprotein-binding protein 1
HMGCR: 3-hydroxy-3-methylglutaryl-CoA reductase
HSPG: heparan sulfate proteoglycan
LD: lipid droplet
LMF1: lipase maturation factor 1
LPL: lipoprotein lipase
LPDS: lipoprotein-depleted serum
PLIN2: perilipin 2
SCD: stearoyl-CoA desaturase
SFM: serum-free media
TG: triglyceride
VLDL: very low-density lipoprotein
VLDLR: very low-density lipoprotein receptor

