## Supplemental Information for "Endocytosis of very low-density lipoprotein particles: an unexpected mechanism for lipid acquisition by breast cancer cells"

### SUPPORTING INFORMATION

**Table S1.** Human cell lines used in these studies and their respective biomarker status and subtype. ER (estrogen receptor), PR (progesterone receptor), HER2 (ErbB2).

| Cell Line | Source | Tumor Type | Subtype | ER | PR | HER2 |
|---|---|---|---|---|---|---|
| BT-474 | Primary breast | Ductal carcinoma | Luminal B | + | + | + |
| DU4475 | Mammary gland | Epithelial cell | TNBC, IM subtype | - | - | - |
| HeLa | Cervix | Adenocarcinoma | N/A | | | |
| MCF-7 | Pleural effusion | Invasive ductal carcinoma | Luminal A | + | + | - |
| MCF10A | Primary breast | Epithelial cell, nontumorigenic | Basal | - | - | - |
| MDA-MB-231 | Pleural effusion | Adenocarcinoma | Claudin-low or basal-like | - | - | - |
| LiSa-2 | Pleomorphic liposarcoma | Liposarcoma | N/A | | + | |
| SKBR3 | Pleural effusion | Adenocarcinoma | Luminal | - | - | + |
| T47-D | Pleural effusion | Invasive ductal carcinoma | Luminal A | + | + | - |

**Table S2.** Gene primer sequences used for detection in qRT-PCR analysis.

| Gene (GenBank) | Product Designation |
|---|---|
| ACACA | Hs01046047_m1 ACACA |
| ACLY | Hs00982738_m1 ACLY |
| ACSS2 | Hs01122829_m1 ACSS2 |
| Cyc | 4310883E |
| CD36 | Hs00354519_m1 CD36 |
| DGAT1 | Hs01020362 g1 |
| FASN | Hs01005622_m1 FASN |
| GPIHBP1 | Hs01564843_m1 GPIHBP1 |
| HMGCR | Hs00168352_m1 HMGCR |
| LDLR | Hs 01092524_m1 LDLR |
| LMF1 | Hs01071616_m1 LMF1 |
| LPL | Hs00173425_m1 LPL |
| PLIN2 | Hs00605340_m1 PLIN2 |
| SCD | Hs01682761_m1 SCD |

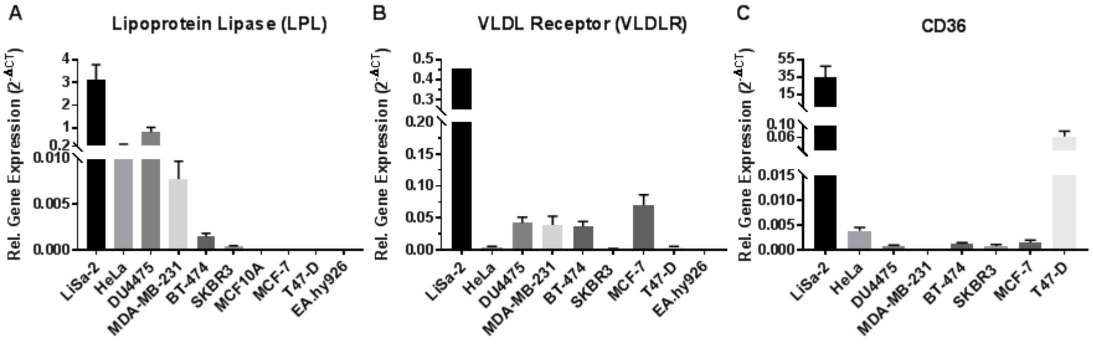


**Figure S1. Cell line expression of select genes involved in lipid uptake.** Basal gene expression of (**A**) LPL, (**B**) VLDLR, and (**C**) CD36 were assessed using qRT-PCR (note different scales). Relative expression values are displayed as $2^{-\Delta CT}$, with Cq of the target normalized to that of cyclophilin. Data are mean +/- SEM of > 3 experiments.

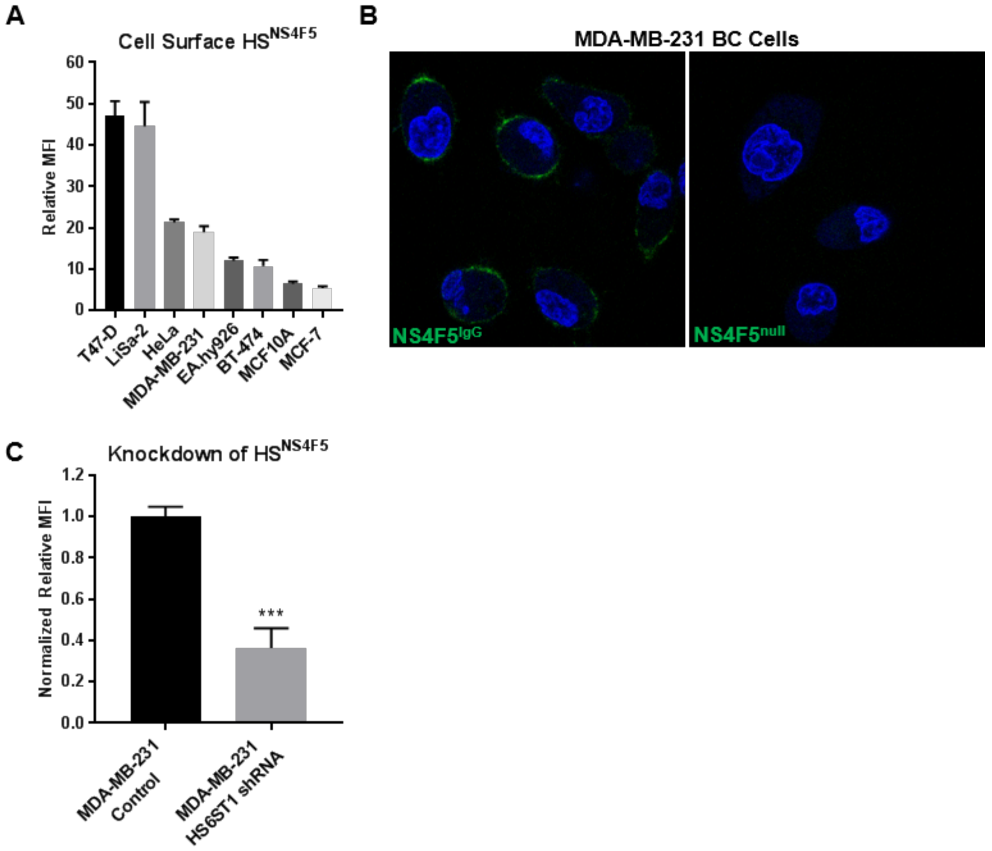


**Figure S2. The HSPG Motif (GlcNS6S-IdoA2S)$_3$, a binding site for LPL, is expressed on the surface of cancer cells.** (**A**) The abundance of HS$^{NS4F5}$ on the surface of cell lines was assessed by flow cytometry. Relative MFI is the median fluorescence of cells stained with the NS4F5$^{IgG}$-Dylight 650 antibody normalized to the NS4F5$^{null}$-Dylight 650 control. (**B**) Confocal microscopy of MDA-MB-231 cells stained with Hoechst 33342 nuclear stain (blue) and DyLight 650 NS4F5$^{IgG}$ or NS4F5$^{null}$antibody (green) for 30 min at 4°C localizes HS$^{NS4F5}$ to the cell surface. (**C**) shRNA knockdown of heparan sulfate 6-O-sulfotransferase 1 (HS6ST1) reduces HS$^{NS4F5}$ on the surface of MDA-MB-231 BC cells, assessed by flow cytometry. Relative MFI represents the median fluorescence of cells stained with the NS4F5$^{IgG}$-Dylight 650 antibody normalized to those stained with the isotype control, NS4F5$^{null}$-Dylight 650. ***$p < 0.001$, two-tailed unpaired t-test with Welch's correction.

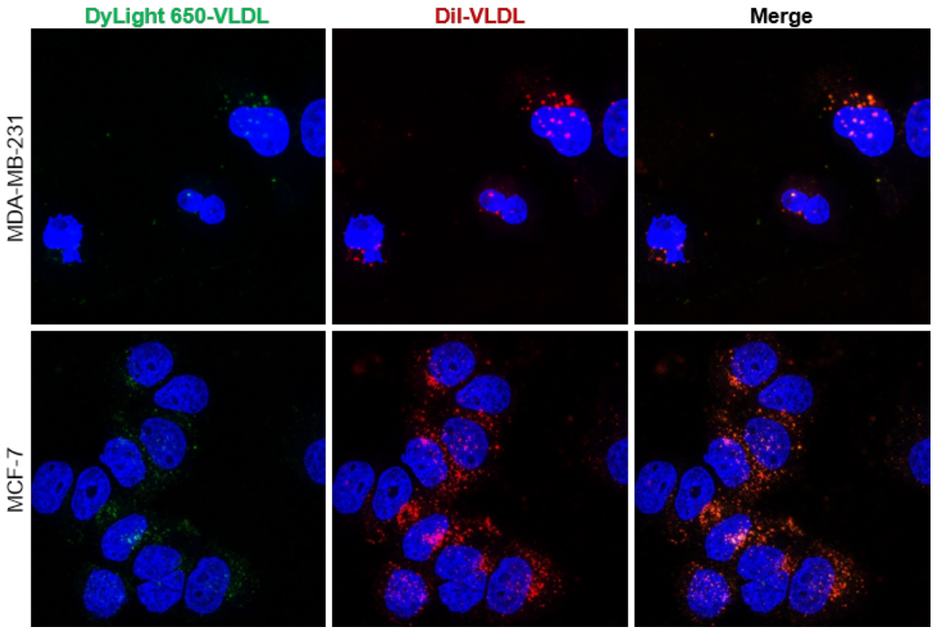


**Figure S3. Both lipid and protein component of VLDL particles bind to- and are internalized by BC cells.** Confocal microscopy of BC cells incubated with DiI (lipid-) and DyLight 650 (protein-) co-labeled VLDLs. Three channels are shown: Hoechst 33342 nuclear stain (blue), DiI-VLDL (red), DyLight650-VLDL (green). White spots (in merged channel) are where DiI and DyLight650 fluorescence coincide.

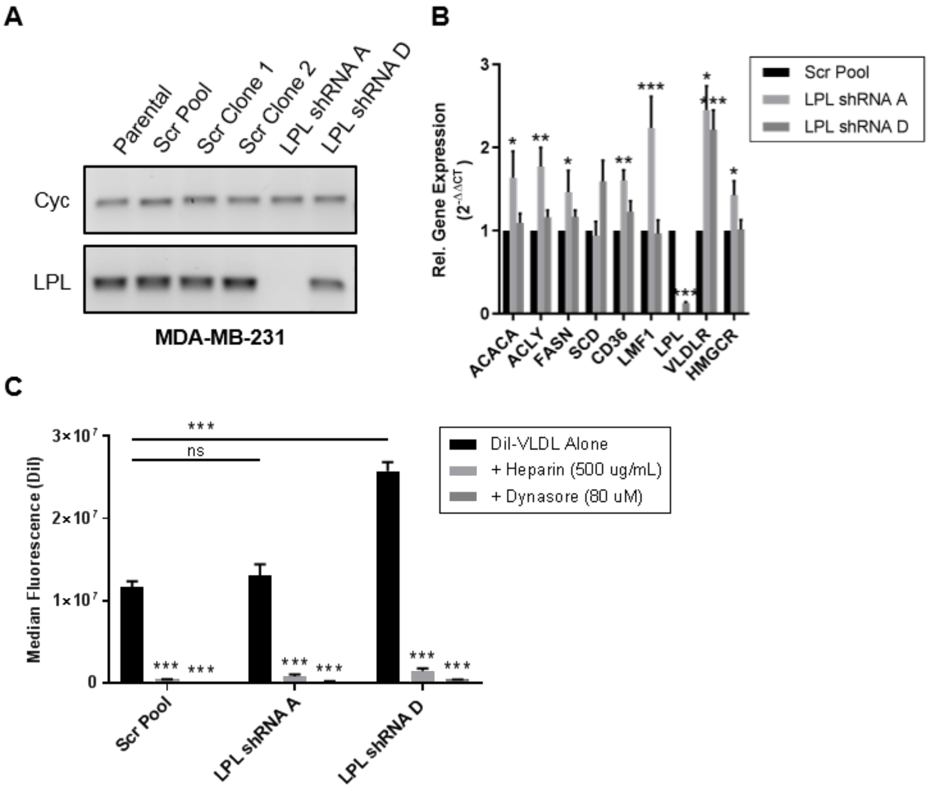


**Figure S4. Characterization of MDA-MB-231 LPL shRNA cell lines.** (**A**) qRT-PCR products run on agarose gels. Parental and scrambled shRNA cell lines are compared to LPL shRNA A (complete knockdown of LPL) and LPL shRNA D (partial knockdown). (**B**) qRT-PCR of MDA-MB-231 manipulated cell lines normalized to the pooled control. LPL shRNA A cells exhibit significant upregulation of mRNA involved in *de novo* lipid synthesis (ACACA, ACLY, FASN), cholesterol synthesis (HMGCR), and FA uptake (CD36, LMF1, VLDLR). MDA-MB-231 LPL shRNA D (partial LPL knockdown) cells display a significant upregulation of VLDLR expression alone. *$P < 0.05$, **$p < 0.01$, ***$p < 0.001$. Error is SEM. (**C**) Comparison of DiI-VLDL uptake in Scr and LPL shRNA cells with and without treatment (heparin 500 µg/mL; dynasore 80 µM), measured by flow cytometry. MDA-MB-231 LPL shRNA D cells had significantly higher DiI-VLDL uptake than either the Scr control or LPL shRNA A cells, ***$p < 0.001$. All three cell lines showed significant reductions in DiI-VLDL uptake following treatment with heparin or dynasore, ***$p < 0.001$.

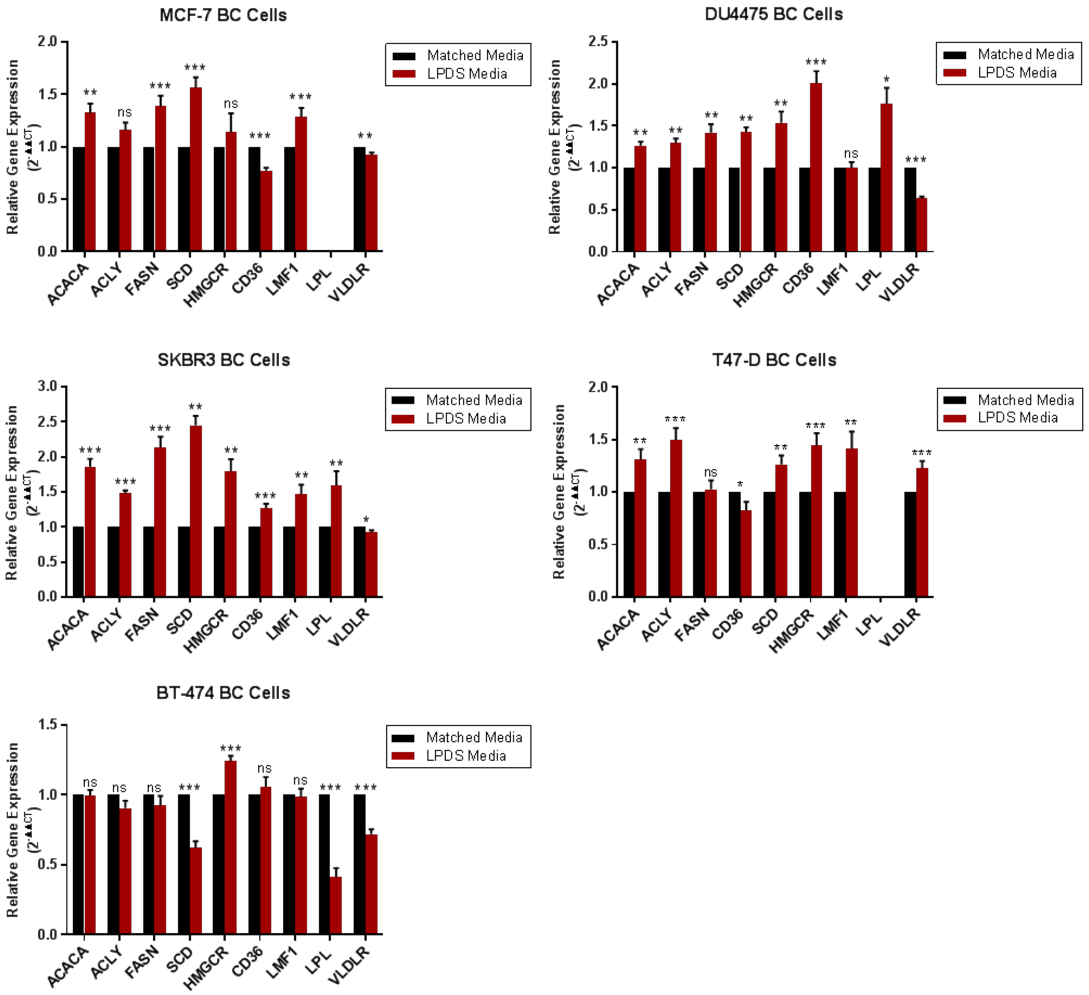


**Figure S5. Trends in the expression of FA metabolic genes in BC cell lines cultured in lipoprotein-depleted serum media for 96 h.** Gene expression was measured using qRT-PCR. Results were calculated using the $2^{-\Delta\Delta CT}$ method first normalized to cyclophilin, and then made relative to that of the matched FBS media control. Data are from > 3 experiments; error is SEM. Statistical significance determined using two-tailed unpaired t-tests: *P < 0.05, **p < 0.01, ***p < 0.001.
